# Structural Lung Remodeling Precedes Functional Decline After Chronic Smoldering Douglas Fir Smoke Exposure in Apoe^-/-^ Mice

**DOI:** 10.64898/2026.05.26.727972

**Authors:** Charbel T. Yazbeck, Jacqueline Matz, Matthew J. Eden, Shubham Rajput, Ye Chen, Michael J. Gollner, Paola Sebastiani, Chiara Bellini, Jessica M. Oakes

**Affiliations:** Department of Bioengineering, Northeastern University, Boston, MA 02115, United States; Tufts Clinical and Translational Science Institute (CTSI), Tufts Medical Center, Boston, MA 02115; Department of Mechanical Engineering, University of California, Berkeley, CA 94720, United States; Institute for Clinical Research and Health Policy Studies (ICRHPS), Tufts Medical Center and Tufts University School of Medicine, Boston, MA 02115

**Keywords:** wildfire smoke, occupational exposure, wildland firefighter, pulmonotoxicology, emphysema, lung mechanics

## Abstract

Wildland firefighters experience repeated exposure to wildfire smoke, yet the pathophysiological mechanisms underlying chronic inhalation injury remain poorly understood. Although prior studies report parenchymal destruction following prolonged woodsmoke exposure, the temporal relationship between molecular, structural, and functional decline following inhalation of smoke from needles/leaves remains unclear. To address this gap, we characterized coordinated changes in lung structure, function, and underlying molecular disruptions using a dosimetry-based murine model approximating 7-14 years of firefighter service. Male apolipoprotein E-deficient mice were exposed to smoldering Douglas fir needle smoke (40 mg/m³, 2 h/day, 5 days/week) for 8 or 16 weeks. Immunofluorescence analyses revealed an early elastolytic response at 8 weeks, with increased neutrophil elastases and matrix metalloproteinases-9 and -12, accompanied by elevated surfactant protein-D, compared to air controls. These changes were resolved by 16 weeks despite progressive tissue injury. Airspace enlargement was evident at 8 weeks, progressed by 16 weeks, and included increased alveolar blunting and septal wall thickening at the later time point. Cleaved caspase-3 was elevated at 16 weeks, indicative of advanced parenchymal damage and apoptosis. Epithelial tight-junction protein ZO-1 intensity was reduced at both evaluation points, whereas the epithelial-to-mesenchymal marker N-cadherin remained undetectable in the alveolar epithelium. Functional impairment as evident by increased static compliance and upward shifts in pressure-volume curves was only significant after 16 weeks of exposure. Findings indicate that molecular and structural injury of tissue destruction preceded measurable functional decline, underscoring the need for early biomarkers to identify smoke-induced lung injury in wildland firefighters before function loss occurs.

## INTRODUCTION

Wildfires have reached an unprecedented level in intensity and frequency, fueled by hotter, drier, and windier climate conditions and the expansion of anthropogenic activities near wildland regions (Andela et al. 2019; Sevinc et al. 2020; Jones et al. 2022). This trend is exemplified by the record-breaking 2021 heat wave in western North America (Thompson et al. 2022; White et al. 2023), the devastating 2023 Canadian wildfires (Kolden et al. 2024), and the recent 2025 Palisades and Eaton fires in California (Babrauskas 2025; Woolcott 2025).

Beyond their direct destructive force, wildfires pose an escalating threat to human respiratory health, primarily due to exposure to wildfire smoke (WFS). A wildfire can release highly concentrated smoke that is composed of a complex mixture of different respirable chemicals, including well known health hazards and air pollutants such as primary fine particulate matter (PM_2.5_), carbon monoxide (CO), nitrogen oxides (NOx), and volatile organic compounds (VOCs) (U.S. EPA 2025). The 2018 Camp Fire was associated with a dramatic increase in PM₂.₅, with concentrations exceeding by more than threefold compared to the levels measured during the same time of year between 2010 and 2017, reaching peaks of 400 µg/m³ (CARB 2021). The recent 2025 Palisades and Eaton Fires resulted in a peak PM_2.5_ -Pb (lead) concentration of 0.5 µg/m³ (Baliaka et al. 2025), nearly a fourfold increase compared to the 2018 Camp Fire.

Rise in wildfire-specific PM_2.5_ has been associated with a more pronounced increase in respiratory hospitalizations than PM_2.5_ from other sources (Aguilera et al. 2021). Globally, more than 300,000 premature deaths annually have been attributed to WFS exposure (Johnston et al. 2012). A growing body of epidemiological evidence links PM_2.5_ exposure from WFS to respiratory and cardiovascular complications, as well as an increase in all-cause mortality (Liu et al. 2015; Reid et al. 2016; Stowell et al. 2024). However, the underlying functional, structural and molecular changes remain inadequately understood.

While the general population experiences significant health risks from WFS exposure, wildland firefighters (WLFFs) face even greater dangers due to their repeated, prolonged and unavoidable exposure during active fire suppression and mop-up efforts, placing them at an elevated risk of respiratory and cardiovascular diseases (Adetona et al. 2016; Black et al. 2017; Navarro et al. 2021). Unlike structural firefighters, WLFFs work in rugged terrain, where self-contained breathing apparatuses are impractical, and they rarely rely on masks beyond a simple bandana. Such coverings provide minimal filtration of PM₂.₅ and do not meet occupational respiratory protection standards required to substantially reduce wildfire smoke inhalation (Adetona et al. 2016; Garg et al. 2023; DHS 2025; Filiberti et al. 2025). The growing intensity of wildfires has also placed greater demands on WLFFs, requiring longer deployments and heavier workloads.

Current epidemiological studies on WLFFs report inconsistent outcomes, with some showing reversible (Gaughan et al. 2008), and others persistent (Jacquin et al. 2011; Panumasvivat et al. 2024) declines in forced expiratory volume in one second (FEV_1_) and forced vital capacity (FVC) post-season. Although the literature on long-term WFS exposure remains sparse, Mathias *et al*. postulate that repeated seasonal inhalation of PM_2.5_ may lead to a sustained decline in spirometry parameters, such as FEV_1_ and FVC (Mathias et al. 2020).

The seasonal nature of WLFF employment, and variations in firefighting jobs, terrain, and exposure duration make it difficult to directly study WLFFs. These challenges also contributes to inconsistencies in wildland firefighter study findings (Ma et al. 2024; Wah et al. 2025). There is a need to provide WLFFs with information regarding the respiratory health consequences due to occupational exposure. Motivated by this gap, we aimed to characterize changes in respiratory structure and function, linked to underlying molecular disruptions, in a controlled laboratory setting. We do this by employing a whole-body murine exposure system designed to model repeated WFS inhalation in WLFFs.

Short-term studies of animal whole-body smoke exposure have been instrumental in characterizing the immediate biological responses that promote acute respiratory events (Migliaccio et al. 2013; Kim et al. 2018; Hargrove et al. 2019; Kim et al. 2019; Ihantola et al. 2020; Ramos et al. 2021; Fiamingo et al. 2024). However, existing long-term animal studies remain sparse and constrained to woodsmoke (Tesfaigzi et al. 2002; Seagrave et al. 2005; Reed et al. 2006; Ramos et al. 2009; Zou et al. 2014; He et al. 2017; Buford et al. 2024; Cochran et al. 2024). In agreement, Ramos et *al.*, Zou *et al*. and He *et al*., all report increased matrix metalloproteinases (MMPs) following woodsmoke exposure, which are key mediators of extracellular matrix degradation (Ramos et al. 2009; Zou et al. 2014; He et al. 2017). When dysregulated, degradation of the alveolar septa drives abnormal enlargement of distal airspaces, a cornerstone of emphysema pathogenesis (Sharafkhaneh et al. 2008). In rats exposed to 1 or 10 mg/m^3^ concentrations of wood smoke for 4 or 12 weeks, there was no detectable change in quasi-static compliance (a measure of lung elastic recoil) (Tesfaigzi et al. 2002). Critically, none of these previous studies directly connected molecular changes to structural and functional impairment. In this study, we fill this gap by relating molecular, structural, and functional changes following prolonged Douglas Fir needle smoke (DFS) exposure.

Our group has developed a mouse inhalation protocol to simulate firefighter exposure under controlled conditions, allowing for precise examination of functional, structural, and molecular changes. Notably, we implemented a whole lung dosimetry framework to relate mouse exposures to a wildland firefighter career duration (Eden et al. 2023; Eden et al. 2025). We equate the cumulative deposited mass of PM, normalized by the lung surface area, between the mouse and the WLFF. We previously modeled 3-7 years of firefighting service and found increased arterial stiffness, reduced left ventricular function, enhanced airway resistance, and increased parenchymal neutrophil elastase and CD68+ cells following 8 weeks of DFS exposure to smoke concentrations of 20 mg/m^3^ (Eden et al. 2023; Matz et al. 2024) in Apolipoprotein E-deficient (Apoe^-/-^) mice. This current study adds to this previous work by exploring the molecular underpinnings of structural and functional changes following ≥ 7 and ≥ 14 years of occupational exposure to wildland fire smoke, providing critical insights into long-term respiratory health risks following prolonged smoke exposure.

## METHODS

A schematic outlining the study assessments is shown in Figure 1.

**Fig. 1.**
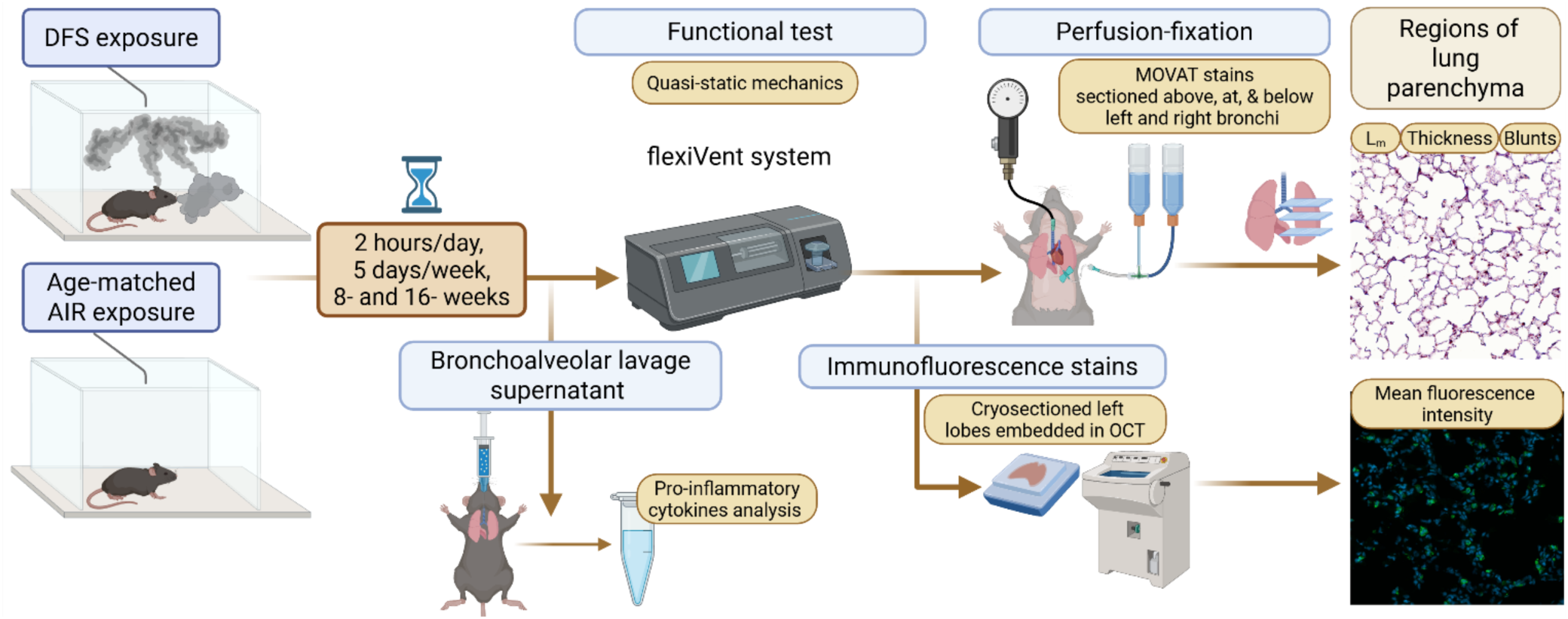
Schematic outlining the functional and structural assessments performed. Following 8- or 16- weeks of filtered air (AIR) or smoldering Douglas fir smoke (DFS) exposure, mice were randomly assigned to experimental groups, including: AIR_8wk_, AIR_16wk_, DFS_8wk_,and DFS_16wk_. Within 48 hours of the final exposure, N = 14, 9, 10, and 10 mice were used for functional assessment in the AIR_8wk_, AIR_16wk_, DFS_8wk_, and DFS_16wk_ groups, respectively. Separate cohorts (N = 5-8 mice per experimental group) were used to assess pro-inflammatory cytokines in bronchoalveolar lavage fluid (BALF) supernatant. Functional assessments included quasi-static measurements (pressure-volume loops, quasi-static compliance). Following functional measurements, lungs were perfusion-fixed or the left lobes were fresh-frozen in optimal cutting temperature (OCT) compound for histological and immunohistochemical analyses, respectively. This was followed by morphological and biomarker assessment of the lung parenchyma. Figure was created with BioRender.com.

### Animals

All animal procedures were approved by the Northeastern University Institutional Animal Care and Use Committee (IACUC) and conducted in accordance with National Institutes of Health (NIH) guidelines. Sexually mature, 8-week-old male Apoe^-/-^ mice, bred on a C57BL/6 background, were obtained from The Jackson Laboratory (Bar Harbor, ME, United States). Apoe knockout mice were used because they exhibit increased cholesterol levels, providing a substrate for oxidative damage and exacerbating the pathophysiological effects of environmental air pollution exposure on the cardiopulmonary system. As observed previously, chronic exposure to cigarette smoke leads to cardiovascular and respiratory impairments in Apoe^-/-^ mice comparable to those observed in habitual smokers (Farra et al. 2021; Matz et al. 2023). In contrast, wild-type mice (C57BL/6) maintain low cholesterol levels and limited respiratory changes, even with long-term cigarette smoke exposure (Knight-Lozano et al. 2002).

Upon arrival, all mice were housed in clear plastic cages in groups of up to five per cage within a temperature-controlled vivarium on a 12-hour light/dark cycle. Food and water were provided ad libitum, with access to standard chow. Mice were randomly assigned to experimental groups, including 8-, 16-weeks of Douglas fir smoke exposure groups (DFS_8wk_ and DFS_16wk_), and an age-matched filtered air-control group (AIR_8wk_ and AIR_16wk_). DFS and AIR groups were acclimated in the animal housing facility for at least 3 days prior to the start of the exposures.

### Smoldering Douglas Fir Smoke Exposure

We previously established a DFS exposure protocol using a dosimetry-based model to match cumulative PM doses between WLFFs and mice, ensuring a physiologically relevant exposure paradigm (Eden et al. 2023; Eden et al. 2025). We determined that 65,000 and 130,000 hours of service could be replicated by exposing male Apoe^-/-^ mice to smoldering DFS at a target concentration of 40 mg/m³ for 2 hours/day, 5 days/week, over 8 and 16 weeks, respectively. Accordingly, 8 and 16 weeks of murine exposure were designed to recapitulate the cumulative particulate inhalation burden corresponding to approximately ≥ 7 and ≥ 14 years of WLFF service, respectively.

DFS was generated using a custom-built tube-furnace exposure system designed in accordance with DIN 53436 standards, ensuring controlled combustion and consistent smoke characteristics (Eden et al. 2023; Garg et al. 2023). Douglas fir needles, sourced from Missoula, MT, were used due to their prevalence in fire-prone ecosystems, availability, ease of storage, and being representative of the Northwest wildfire scenarios. Needles were selected rather than wood as they represent the primary fine fuel component that drives wildland fire ignition and spread. Moreover, needle combustion generates a broader and more abundant volatile organic compound emission profile compared to wood combustion (Jain et al. 2012; Pallozzi et al. 2018), thereby providing a more toxicologically relevant exposure model.

Fuels were dried in a 75°C oven for 72 hours, then refrigerated for long time storage with a desiccant pack. For smoldering smoke generation, fuels were loaded inside the quartz tube and the furnace was set to a temperature of 450 °C at a speed of 20 mm/min with a primary air flow rate of 3 L/min. Downstream air flow was added to the system to dilute the smoke to the targeted PM concentration of 40 mg/m^3^ prior to entering the 3D-printed custom whole-body exposure chamber. Particle concentration, CO concentration, temperature, and humidity were monitored throughout the exposure.

### Assessment of respiratory mechanics

Respiratory mechanics were collected within 48 hours of the last day of exposure. Mice (N = 14, 9, 10, and 10 for AIR_8wk_, AIR_16wk_, DFS_8wk_ and DFS_16wk_, respectively) were anesthetized via intraperitoneal injection of a cocktail containing xylazine (20 mg/kg), ketamine (100 mg/kg), and acepromazine maleate (3 mg/kg). Once an appropriate level of sedation was established, mice underwent tracheostomy with a rigid 18-gauge cannula. To prevent spontaneous breathing during the procedure, mice were paralyzed with 30 mg/kg pancuronium bromide after they were connected to the ventilator. Heart rate was continuously monitored using the PhysioSuite Mouse Heart Rate Monitor (Kent Scientific Corp., Torrington, CT, USA) and thermal support was provided.

Quasi-static pressure-volume (P-V, FlexiVent, FX Module 2, SCIREQ, Montreal, QC, Canada with the FlexiWare v.8.0 software) curves were collected using eight incremental pressure steps, beginning at 3 cmH₂O and progressing to 30 cmH₂O, near total lung capacity (TLC). Lungs were then deflated similarly. Calculation of quasi-static compliance (C_st_) was derived from the Salazar-Knowles (Salazar and Knowles 1964) model rather than from the slope at the inflection point of the expiratory limb of the sigmoidal fits, due to high variability in the latter approach.

### Morphometry analysis

Following completion of lung mechanics testing, the lungs were maintained at a constant tracheal pressure of 20 cmH₂O and transcardially perfused (20 cmH₂O) with Ringer’s solution (120 mM NaCl, 5.0 mM KCl, 0.15 mM CaCl₂). This was followed by 3% glutaraldehyde to preserve structural integrity for histological staining. The entire lung block, including lobes and trachea, was extracted and post-fixed at 4 °C for at least 24 hours before being transferred to 70% ethanol for long term storage. The left lung and right apical lobe from N=6 mice per group were selected for morphometric evaluation. These were dehydrated, sectioned transversely above, at, and below the main bronchus and embedded in paraffin.

Thin sections (5 µm) were stained using Movat’s pentachrome stain to visualize structural compartments. High-resolution imaging at 40x magnification was performed using an automated slide scanner (EasyScan Infinity, Motic), and morphometric parameters: alveolar septal thickness and linear mean intercept (L_m_) were extracted using custom MATLAB scripts (Matz et al. 2023). L_m_ quantifies airspace size and is widely adopted to confirm the presence of emphysematous lesions (Fisk and Kuhn 1976; Mercer and Crapo 1992; Vlahovic et al. 1999; Robbesom et al. 2003; Foronjy et al. 2006). In addition, enlarged alveolar spaces can be identified by the presence of the alveolar walls blunts, the so called “drummer stick” (Murărescu et al. 2008). Alveolar blunts were counted by a blinded observer as remnant septal fragments are present in destructed alveolar spaces.

### Immunofluorescence imaging and analysis

Following lung mechanics assessment, N=3-5 freshly isolated left lobes were transversely sectioned each at the main bronchus and both preserved in optimal cutting temperature (OCT) blocks. Three cross-sections (7-10 µm thick) were cryosectioned at - 20°C using a Leica CM3050 cryostat (Leica Microsystems, Wetzlar, Germany) onto positively charged Superfrost Plus microscope slides and stored at -80°C until immunostaining.

Immunofluorescence staining was achieved using standardized protocols. If the primary antibody was derived from a mouse, a mouse-on-mouse blocking reagent (Thermo Scientific, R37621) was used at 1x for 1 h to block endogenous signal interference. The following primary antibodies were used: anti-ZO-1 (339100, Invitrogen, 1:100), anti-N-CAD (AB18203, Abcam, 1:100), anti-MMP9 (29022, QED Bioscience,1:200), anti-MMP12 (22989, Protintech, 1:200), anti-surfactant protein D (AB220422, Abcam, 1:200), anti-neutrophil elastase (115648, Invitrogen, 1:200), anti-cleaved caspase-3 (9664s, Cell Signaling Technology, 1:1000), anti-cleaved caspase-9 (9509, Cell Signaling, 1:100), anti-NF-κB p65(AB16502, Abcam, 1:100), anti-phospho- NF-κB p65 (3033, Cell Signaling, 1:500). For nuclear counterstaining, the samples were incubated with DAPI (62248, Thermo Scientific, 1:5000) diluted in 1x PBS for 10 min. Slides were then mounted using Epredia™ Immuno-Mount™ (9990402, Thermo Scientific) and sealed with nail polish prior to imaging.

For each mouse, at least one random parenchymal region was imaged in each of the three lung cross sections per slide. Regions containing airways or vasculature were excluded from parenchymal assessments. Negative control slides, in which the primary antibody was replaced with 1x PBS, were included in each staining experiment to confirm the antibody specificity. Imaging was performed using a Zeiss LSM 800 confocal microscope at 20x or 40x magnification. Consistent acquisition parameters were maintained across all samples for each marker using Zeiss Efficient Navigation (ZEN) software (Carl Zeiss Microscopy, Jena, Germany). Images were saved in CZI format and imported as grayscale split channels using the “Bio-Formats Import Options” in FIJI (NIH, Bethesda, MD, USA) for analysis.

For image quantification, the measurement of mean fluorescence intensity per tissue surface area was automated using a custom macro script. Briefly, images were converted to grayscale and then the region of interest (ROI), i.e. tissue layer, was then masked by segmentation of tissue autofluorescence using the thresholding method. The intensity was measured within the borders of the masked region after background subtraction using an autofluorescence threshold. NF-κb p65 nuclear intensity was measured for every nucleus by segmenting DAPI and then averaging the sum of nucleic signal by the total number of cells per image. Each mask was saved and visually inspected and confirmed for accuracy and overlap with the region of interest.

### Statistical analysis

We conduct group-level comparisons using generalized linear models (GLMs). We employed Analysis of variance (ANOVA) to evaluate main effects and to test interaction effects between factors. For comparisons of longitudinal trends, we assessed pointwise differences between groups using ANOVA at each time point. We evaluated overall trajectory differences using mixed-effects models to account for within-subject correlation over time. For analysis involving nested or clustered data structures—such as repeated measures where multiple images were taken for the same section, we used mixed-effects models with appropriate random effects to account for within-cluster correlation. We also calculated Intraclass correlation coefficients (ICCs) to quantify the proportion of variance attributable to clustering and to evaluate the degree of within-cluster dependence. To control multiple hypothesis testing, we adjusted p-values using the false discovery rate (FDR) method, applying an FDR threshold of 0.10. Statistical significance was evaluated using two-sided tests, with adjusted p-values reported where applicable.

All analysis was conducted using R version 4.3.0.

## RESULTS

The exposure system delivered stable and respirable smoke with a particle count median diameter of 110 ± 20 nm (GSD 1.47 ± 0.03) (Eden et al. 2023), a particulate mass concentration of 39 ± 13 mg/m³, and a CO concentration of 218 ± 46 ppm. Exposure chamber temperature was 23.7 ± 1.1 °C and relative humidity was 42.8 ± 11.2%. Under these exposure conditions, DFS-exposed mice exhibited a dampened increase in percent initial body mass compared to AIR-exposed mice at the 8 (AIR: 22%, DFS: 18%) and 16 (AIR: 19%, DFS: 12%) weeks exposure durations.

### DFS exposure triggers early elastolytic response

Growing evidence suggests that repeated and prolonged woodsmoke exposures induce emphysematous changes in animal lungs (Ramos et al. 2009; Zou et al. 2014; He et al. 2017), a form of chronic obstructive pulmonary disease (COPD). To explore this, we immunofluorescence stained for markers responsible for elastin degradation (elastolysis): neutrophil elastase (NE), MMP-12, and MMP-9 (**Error! Reference source not found.**) (Finlay et al. 1997; Ohnishi et al. 1998; Imai et al. 2001; Russell et al. 2002; Molet et al. 2005; Demedts et al. 2006 Mar 1; Demkow and van Overveld 2010; McGarry Houghton 2015; Gramegna et al. 2017). Elastin is a load-bearing component of the alveolar extracellular matrix and maintains elastic recoil. Its degradation compromises structural integrity, often leading to alveolar wall rupture. The quantity of positively stained markers of NE, MMP-12, and MMP-9 was higher in the DFS than AIR exposed mice at 8-week (FDR<0.0001, Fig. 2B; FDR = 0.0029, Fig. 2C; FDR =0.0199, Fig. 2D), but not at the 16-week timepoint.

**Fig. 2.**
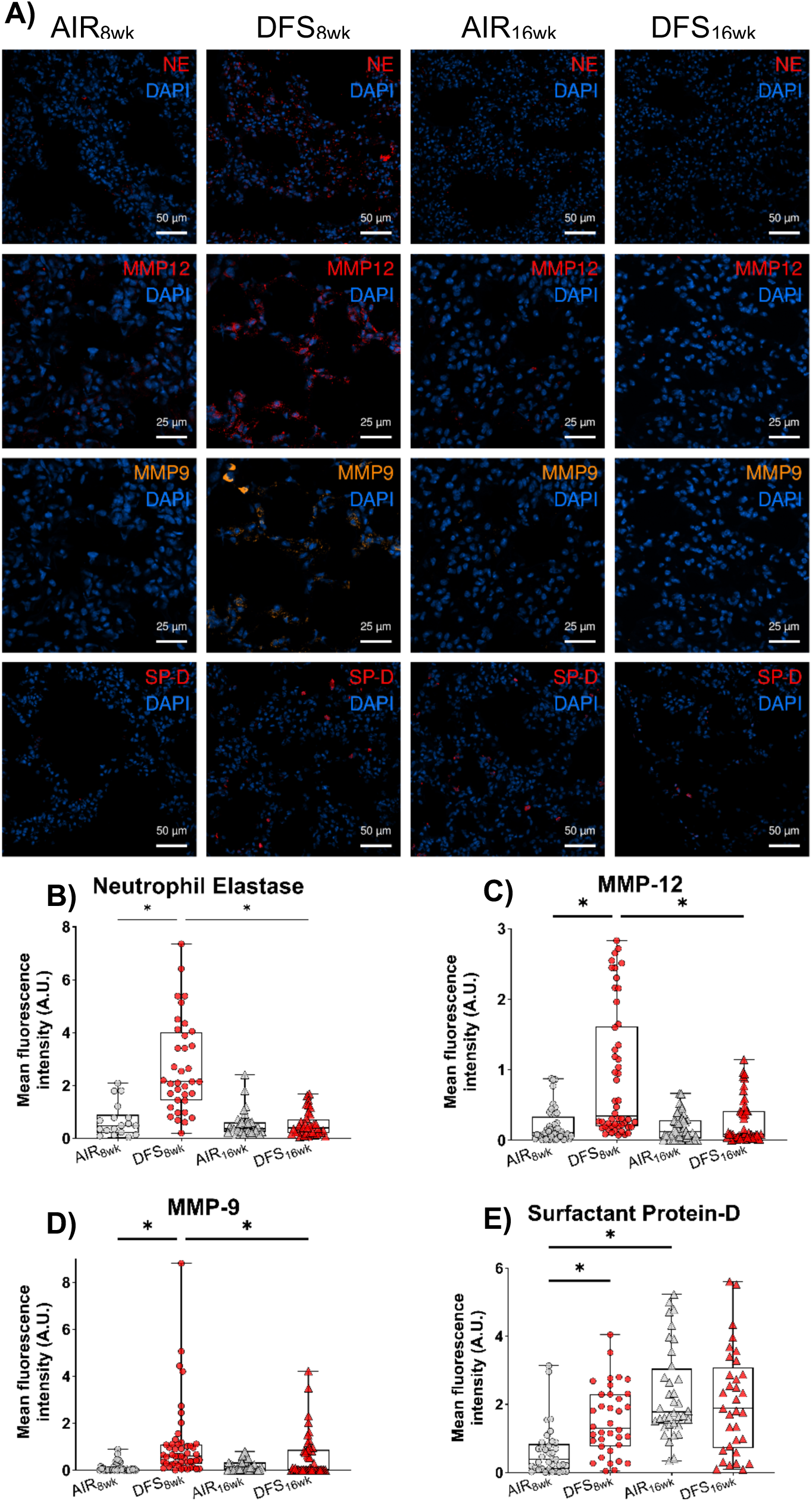
Immunofluorescent analysis of elastolytic enzymes and surfactant protein D in the lung parenchyma following DFS exposure. (A) Representative confocal images showing neutrophil elastase (NE), macrophage metalloelastase (MMP-12), matrix metalloproteinase-9 (MMP-9), and surfactant protein D (SP-D) expression of random parenchymal regions from AIR_8wk_, DFS_8wk_, AIR_16wk_, and DFS_16wk_ groups. MMP-12 (red) and MMP-9 (orange) were double-stained, while NE, SP-D (red) were stained independently. DAPI nuclear staining (blue) was included in all images. Quantification of normalized fluorescence intensity per tissue surface for (B) NE, (C) MMP-12, (D) MMP-9, and (E) SP-D in alveolar regions of the lung, expressed in arbitrary fluorescence units (A.U.) showed a transient pattern, with a peak of all markers for the DFS_8wk_ condition. Images were captured under the same acquisition parameters and magnification per marker. Magnifications of 20x or 40x were used; scale bars are shown at 50 μm or 25 μm, respectively. Individual data points represent mean fluorescence intensity per image. Box plots show the median and interquartile range, with whiskers indicating minimum and maximum values (N = 3-4 mice/group; multiple cross-sections and parenchymal regions per lung). Immunofluorescence data were analyzed using mixed-effects models accounting for within-mouse clustering (ICC), with pairwise comparisons adjusted using FDR; significance was defined for FDR-adjusted p < 0.1 (*).

Surfactant protein-D (SP-D) is an emerging COPD marker (Bowler 2012) and it is an immune-modulatory surfactant protein produced by alveolar type II cells. It has been shown to guard against emphysema, in particular, by preventing secretion of MMP-9 and MMP-12 (Wert et al. 2000; Yoshida and Whitsett 2006; Pastva et al. 2007; Watson et al. 2021; Shamim et al. 2024). Similar to the pattern observed with elastolytic enzymes, SP-D fluorescence intensity also exhibited a transient response in the DFS-exposed group. SP-D fluorescence intensity was significantly higher in DFS_8wk_ than AIR_8wk_ (FDR = 0.0505, **Error! Reference source not found.**E). SP-D fluorescence intensity became more prominent with longer air exposure: AIR_16wk_ significantly increased compared to AIR_8wk_ (FDR = 0.0001, **Error! Reference source not found.**E), while longer DFS exposure did not change SP-D fluorescence. This resulted in no difference between groups at the 16-week time point.

### DFS exposure causes morphometry changes in lung parenchyma

Emphysema is characterized by the irreversible and progressive degradation of alveolar walls and enlargement of distal airspaces, fueled by irregulated elastolysis. The sizes of airspaces were the largest for the DFS_16wk_ group, as evidenced qualitatively by visible signs of emphysematous lesions (**Error! Reference source not found.**D), by the largest L_m_ (**Error! Reference source not found.** and an elevated number of alveolar blunts (**Error! Reference source not found.**F, FDR = 0.04 compared to AIR_16wk_). In addition, L_m_ was significantly lower in AIR_8wk_ but greater in DFS_16wk_ than DFS_8wk_, respectively (FDR = 0.0449, FDR= 0.0036 respectively), indicating progressive airspace enlargement with prolonged DFS exposure. Moreover, AIR_16wk_ mice had significantly larger airspaces than AIR_8wk_ mice (FDR = 0.0449), indicating an aging effect in Apoe^-/-^ mice (Massaro and Massaro 2011).

Structurally, parenchymal septal walls thickened with DFS exposure (**Error! Reference source not found.**G) at both time points compared to matched AIR controls (FDR<0.0001 and FDR = 0.0002 for 8- and 16-week time points). In contrast, parenchymal walls thickness decreased with continued DFS exposure and age, thickness was significantly larger in DFS_8wk_ than DFS_16wk_ (FDR=0.0012) and in AIR_8wk_ than AIR_16wk_ (FDR = 0.0481), respectively.

### DFS exposure compromises lung integrity

NF-κb p65 is a smoke-responsive master mediator of inflammation and epithelial-to-mesenchymal transition (EMT), underscoring septal and airway tissue remodeling (Zhang et al. 2016; Tian et al. 2017; Markopoulos et al. 2019; Chen et al. 2020) in inhalation toxicology such as ambient PMs (Chi et al. 2018; Xu et al. 2019; Chen et al. 2020), and cigarette smoke (Zhao et al. 2013; Lu et al. 2018; Ma et al. 2020; Zhou et al. 2020; Su et al. 2022). To interrogate EMT, we conducted dual immunostaining for epithelial and mesenchymal markers, Zonula Occludens-1 (ZO-1) and N-Cadherin (N-Cad), respectively (**Error! Reference source not found.**A, B).

There was a significant reduction in ZO-1 intensity in the alveolar epithelium of DFS-exposed mice at both the 8-week and 16-week timepoints, compared to age-matched AIR-exposed controls (FDR = 0.0995, FDR = 0.0616, **Error! Reference source not found.**A, B, respectively). N-Cadherin expression remained undetectable in the parenchyma across all experimental groups. N-Cadherin positive signal was observed and confirmed in vascular endothelium, consistent with prior reports (Gentil-dit-Maurin et al. 2010). Reduced ZO-1 intensity in the absence of N-Cadherin indicates barrier weakening independent of complete EMT.

To investigate this further, we stained for NF-κB p65 and found that it did not localize to the nucleus (Suppl, Fig. S2). Pro-inflammatory cytokines levels also were not significantly different (Suppl, Fig. S1) in the collected bronchoalveolar lavage fluid of DFS smoke exposed mice to control.

### DFS exposure triggers apoptotic response

Programmed cell death (apoptosis) is another process implicated in emphysema pathogenesis, even in the absence of overt inflammation (Kasahara et al. 2000; Aoshiba et al. 2003; Tang et al. 2004; Petrache et al. 2005; Demedts et al. 2006; Fehrenbach 2007). Cleaved caspase-3 (CC-3) was selected due to its strong indication for cells undergoing irreversible apoptosis (Duan et al. 2003; Glamočlija et al. 2005). Parenchymal CC-3 intensity was higher in DFS than AIR at 16-week (FDR = 0.0003, **Error! Reference source not found.**A), but not at the 8-week timepoint. When CC-3 staining was observed in the nucleus, there was consistent co-localization with distinct nuclear condensation and fragmentation, hallmarks of late-stage apoptosis (Häcker 2000), that were present in the DFS_16wk_ group (**Error! Reference source not found.**A). This was further supported by a similar pattern in its upstream effector, cleaved caspase-9 (CC-9, Supplementary Material, Fig. S3) (Porter and Jänicke 1999; Stegh and Peter 2001). While CC-9 positive intensities were observed across all groups, its intensity was consistently the highest in DFS_16wk_ (FDR =0.08 compared to AIR_16wk_).

### DFS exposure impairs lung function

The quantitative assessment of lung function through ventilator-assisted techniques has long been considered a gold standard in animal models of emphysema (Sterk 2006). The well-established signature of an emphysematous lung is marked with an upward shift of the P-V curve and an associated increase in static lung compliance (MacKlem and Becklake 1963). We focused on pressure-volume (P-V) curves and static compliance (C_st_), widely used parameters for characterizing disease signatures. P-V curves were slightly elevated in DFS_8wk_ compared to AIR_8wk_ (**Error! Reference source not found.**A), and further increased in DFS_16wk_ relative to AIR_16wk_ (**Error! Reference source not found.**B). Static compliance (C_st_, **Error! Reference source not found.**C), derived from the Salazar-Knowles model fit to the expiratory pressure-volume curve (Salazar and Knowles 1964), was significantly higher in DFS_16wk_ compared to AIR_16wk_ (FDR=0.0697). C_st_ was slightly higher in DFS_8wk_ than AIR_8wk_ without reaching statistical significance (FDR=0.1553). This outlines a decoupling of the structure-function relationship at an early stage, where structural changes precede measurable functional changes.

## DISCUSSION

Extreme fire weather conditions have increased by approximately 65% over the last four decades (1980-2023), with predictions indicating that extreme wildfire events may increase by 30% by 2050 (Abatzoglou et al. 2021; United Nations Environment Programme and GRID-Arendal 2022; Wasserman and Mueller 2023; Cunningham et al. 2025; Matteo et al. 2025). Smoke originating from smoldering biomass is a significant concern as it contributes significantly to smoke-related air pollution (Jaffe et al. 2020; Rein and Huang 2021). We examined whether chronic exposure to DFS perturbs the structure-function relationship of the lungs of Apoe^⁻/⁻^ mice. This study enhances prior chronic biomass smoke studies (Ramos et al. 2009; He et al. 2017), which reported substantial enlargement of the airspaces and elevated apoptosis. We add to this body of work by also quantifying septal thickening and the connection between lung structure and function. We discovered that early molecular and structural emphysematous-like changes precede measurable changes in pulmonary mechanics.

Elastolysis is a process that refers to the degradation of elastin fibers by elastase enzymes such as neutrophil elastase, macrophage metalloelastase (MMP-12), and MMP-9 (gelatinase B). These enzymes initiate the matrix breakdown underlying emphysematous remodeling observed in the lungs of COPD patients (Finlay et al. 1997; Imai et al. 2001; Russell et al. 2002; Molet et al. 2005; Demedts et al. 2006 Mar 1) and robust animal models of emphysema (Hautamaki et al. 1997; Shapiro et al. 2003; Pemberton et al. 2005). Converging lines of evidence from animal models, human studies, and mechanistic experiments highlight a possibility of lung injury via “protease-antiprotease imbalance”, central to emphysema pathogenesis (Demkow and van Overveld 2010; Sandhaus and Turino 2013; Gramegna et al. 2017). For example, mice deficient in MMP-12 (Hautamaki et al. 1997) or neutrophil elastase (Shapiro et al. 2003) are protected from chronic cigarette smoke-induced emphysema. Notably, the same group of researchers demonstrated that the combined action of these two elastases establishes a self-amplifying cycle of tissue degradation. Mechanistically, MMP-12 and NE potentiate one another by inactivating their respective endogenous inhibitors: alpha-1 antitrypsin (A1AT) and tissue inhibitor of metalloproteinases-1 (TIMP-1) in the tissue matrix (Gronski et al. 1997; Shapiro et al. 2003; Demkow and van Overveld 2010). The simultaneous inactivation of extracellular A1AT and TIMP-1 might disrupt protease inhibition and clearance, enabling accumulation of MMP-12 and NE in the matrix, even after exposure ends (Babusyte et al. 2007; Louhelainen et al. 2010). In their study, D’Armiento *et al*. failed to find a real-time association between measured MMP-12 and MMP-9 levels and functional decline in COPD patients and smokers (D’Armiento et al. 2013). In our work, the elasolytic surge preceded a detectable functional decline. Despite resolution of intracellular elastolytic enzyme activity after 16 weeks of DFS exposure, we observed aggravation of tissue destruction leading toward functional decline. In support of this observation, Ramos *et al*. found a temporal separation between peak protease activity and ongoing enlargement of airspaces in guinea pigs exposed to pine wood smoke for up to 7 months (Ramos et al. 2009). Reports in humans (Montaño et al. 2004; Guarnieri et al. 2014) and animal models (Ramos et al. 2009; Zou et al. 2014) exposed to long-term wood smoke also document elevations in MMP-12 and MMP-9, consistent with our early proteolytic trigger for later structural changes.

The stress redistribution-mechanical failure phenomenon, wherein the persistence of cyclic mechanical stress, such as that from the normal breathing cycle, may slowly propagate tissue damage after an initial surge in proteolytic degradation (Kononov et al. 2001; Suki et al. 2003; Suki et al. 2012; Tanabe et al. 2020). This may provide a mechanical explanation for the progression of airspaces enlargement leading to increased lung compliance in the absence of overt inflammation and elastolytic activity. Surfactant protein-D (SP-D) also exhibited a transient pattern that aligned with the elastase dynamics. Synthesized by alveolar type II cells, SP-D has emerged as a key endogenous regulator of immune responses and helps to prevent emphysema. Mice deficient in SP-D develop spontaneous emphysema marked by elevated MMP-9 and MMP-12 expression, which worsens with age (Wert et al. 2000). Under homeostatic conditions, SP-D suppresses macrophage activation by binding to their surface receptors and attenuating the inflammatory response. It does that by regulating NF-κb in alveolar macrophages, a master transcription factor in inflammatory tissue remodeling that drives the expression of pro-inflammatory mediators (Yoshida and Whitsett 2006). Following DFS exposure, we did not observe an activation of NF-κB p65, or detect a pronounced presence of pro-inflammatory cytokines in BALF at our two evaluation points. Although interleukin (IL)-1α and IL-1β expression exhibited a transient pattern, these changes did not reach statistical significance at the intermediate time point. Using a human-controlled woodsmoke exposure designed to replicate wildland firefighting conditions, Ferguson *et al*. reported increased SP-D levels in exhaled breath condensate and systemic circulation following acute exposure (Ferguson et al. 2016). Further investigation is needed to determine if the elevation of SP-D directly contributed to the decline in elastolytic enzymes observed between 8 and 16 weeks of DFS exposure.

In line with our observations, McCarthy *et al*. (McCarthy et al. 2016; McCarthy et al. 2017) failed to observe NF-κB activation in cultured human small airway epithelial cells exposed to 45 or 55 mg/m^3^ smoke for up to an hour at the air-liquid interface (ALI), despite clear evidence of an elastolytic response (Mehra et al. 2012). Instead, they implicated AP-1 signaling in mediating smoke-induced responses. Following this, Mehra *et al*. showed that HSEACs did not express TIMP-1, and therefore concluded that HSAECs fail to counterbalance the biomass smoke-induced elevation of MMP-9 and MMP-12 (Mehra et al. 2012). Similarly, Upadhyay *et al*., showed that NF-κB p65 and pro-inflammatory cytokines levels were not induced following woodsmoke exposure of alveolar epithelial cell line NCI-H441 at ALI after five exposures in three days (Upadhyay et al. 2024). Primary evidence from *in vitro* studies, showed the potential of irritated pulmonary epithelial cells to secrete MMP-9 and MMP-12 (Yu et al. 2010; Morales-Bárcenas et al. 2023). Alongside macrophages and neutrophils, the role of alveolar epithelial cells in contributing to the protease-antiprotease imbalance found during wildfire smoke-induced injury needs further investigation.

Given the established role of NF-κB as a master transcriptional regulator of EMT-driven pulmonary fibrosis following environmental exposures (Tian et al. 2017; Tian et al. 2018; Zhang et al. 2019; Chen et al. 2020; Zhang et al. 2022; Ding et al. 2024), the absence of its activation is consistent with the limited EMT marker induction of N-Cad, observed in our current study. In a study (Zou et al. 2014), where rats were exposed to 27.6 ± 12.1 mg/m^3^ of smoldering China fir sawdust smoke for 4 months, there was no indication of the profibrotic markers FSP1, vimentin, and α-SMA in small airways. Nevertheless, the decrease in ZO-1 intensity following exposure suggests impaired epithelial barrier integrity in the absence of classical EMT-mediated remodeling. In this context, converging evidences support alternative mechanisms of epithelial tight-junction disruption, including direct cleavage of the junctional proteins by NE and MMPs (Peterson et al. 1995; Miyazaki et al. 1998; Bojarski et al. 2004; Boxio et al. 2016), as well as fragmentation of ZO-1 by apoptotic caspases such as CC-3 (Bojarski et al. 2004), collectively compromising epithelial barrier integrity. Consistent with these mechanisms, DFS exposure resulted in sustained reductions in ZO-1 immunostaining across both timepoints (Fig. 4), with elevated elastase markers evident after 8 weeks (Fig. 3) and increased CC-3 (Fig. 5) intensity detected after 16 weeks of DFS exposure. This supports the long-standing hypothesis that ultrafine smoke PMs, such as those in WFS, compromise the air-blood barrier and translocate into the systemic circulation, reaching and impacting extrapulmonary organs (Nemmar et al. 2001; Nemmar et al. 2002; Shimada et al. 2006; Williams et al. 2024). However, this has not yet been studied for wildland fire smoke in an *in vivo* model and warrants further investigation.

**Fig. 3.**
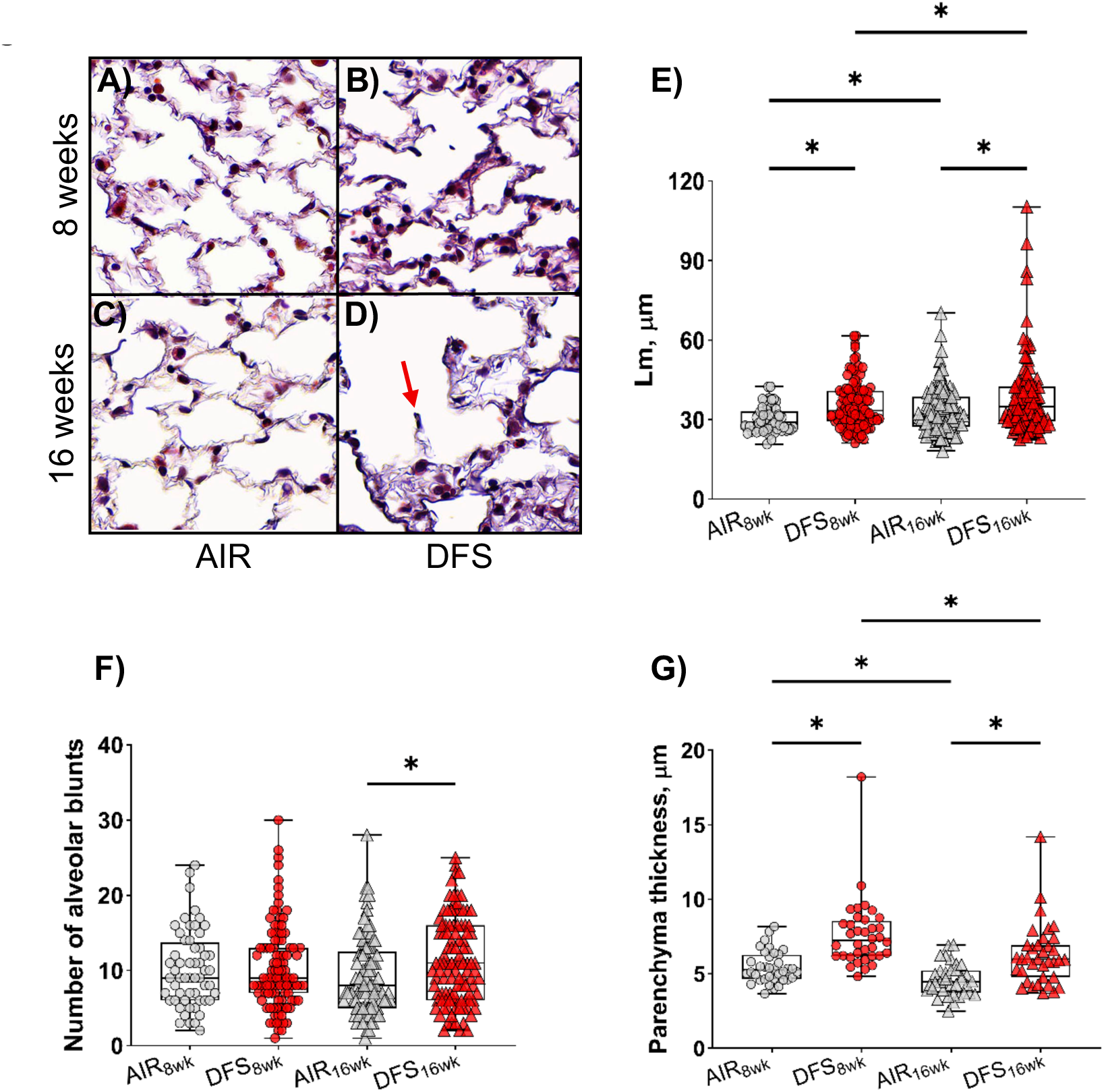
Histological evidence and morphometric quantification of alveolar airspace changes following DFS exposure. Representative MOVAT stained lung sections from A) AIR_8wk_, B) DFS_8wk_, C) AIR_16wk_, and D) DFS_16wk_ groups demonstrate progressive emphysematous lesions, particularly at 16 weeks of DFS exposure. E) Mean linear intercept (L_m_) measurements show a significant increase in alveolar airspace dimensions in both DFS-exposed groups compared to their respective AIR controls. Additionally, L_m_ was significantly elevated in DFS_16wk_ compared to DFS_8wk_, indicating a progressive worsening of alveolar architecture over continued exposure time. F) Alveolar blunt (red arrow in D) counted per each field of view were significantly elevated in DFS_16wk_ compared to AIR_16wk_. G) Parenchyma septal thickness increased with DFS compared to matched AIR controls and decreased with prolonged exposure DFS_16wk_ vs. DFS_8wk_. All images were taken under the same magnification. Data represent the interquartile range shown as boxes, with the middle line indicating the median and whiskers extending to the minimum and maximum values (N = 6 mice per group, with multiple cross-sections analyzed per mouse and multiple parenchymal regions were quantified per cross-section). Group differences were assessed using one-way ANOVA with FDR-adjusted p-values. Statistical significance between experimental groups was defined for FDR-adjusted p < 0.1 (*).

**Fig. 4.**
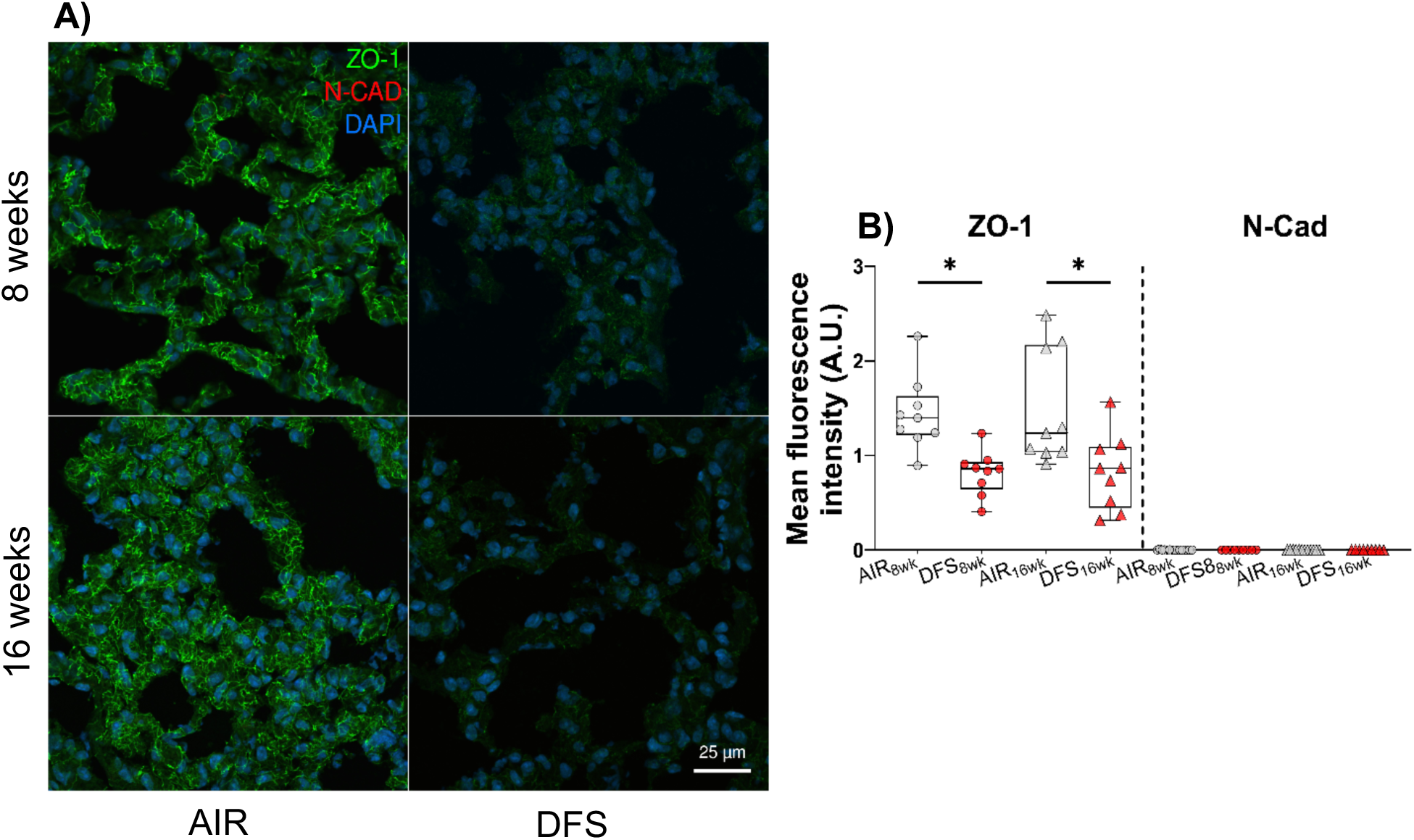
Dual immunofluorescence analysis of epithelial junctional proteins ZO-1 and N-cadherin in the lung parenchyma following DFS exposure. A) Representative confocal images of random parenchymal regions from AIR_8wk_, DFS_8wk_, AIR_16wk_, and DFS_16wk_ groups, stained for the tight junction protein, Zonula Occludens-1 (ZO-1, green) and the adherens junction marker N-Cadherin (N-CAD, red), with DAPI nuclear counterstain (blue). ZO-1 localized to alveolar septa in AIR control lungs, while its intensity was markedly diminished in DFS-exposed lungs. N-Cadherin expression was undetected across all groups. B) Quantification of mean fluorescence intensity of ZO-1 and N-Cadherin in the alveolar region, expressed in arbitrary units (A.U.). ZO-1 signal was significantly reduced in DFS_8wk_ and DFS_16wk_ groups compared to their respective age-matched AIR controls. N-Cadherin levels were uniformly low and unchanged across groups. All images were acquired at 40x magnification and under the same acquisition parameters. Scale bar = 25 μm. Individual data points represent mean fluorescence intensity per image. Box plots show the median and interquartile range, with whiskers indicating minimum and maximum values (N = 3 mice/group; multiple cross-sections and parenchymal regions per lung). Immunofluorescence data were analyzed using mixed-effects models accounting for within-mouse clustering (ICC), with pairwise comparisons adjusted using FDR; significance was defined for FDR-adjusted p < 0.1 (*).

**Fig. 5.**
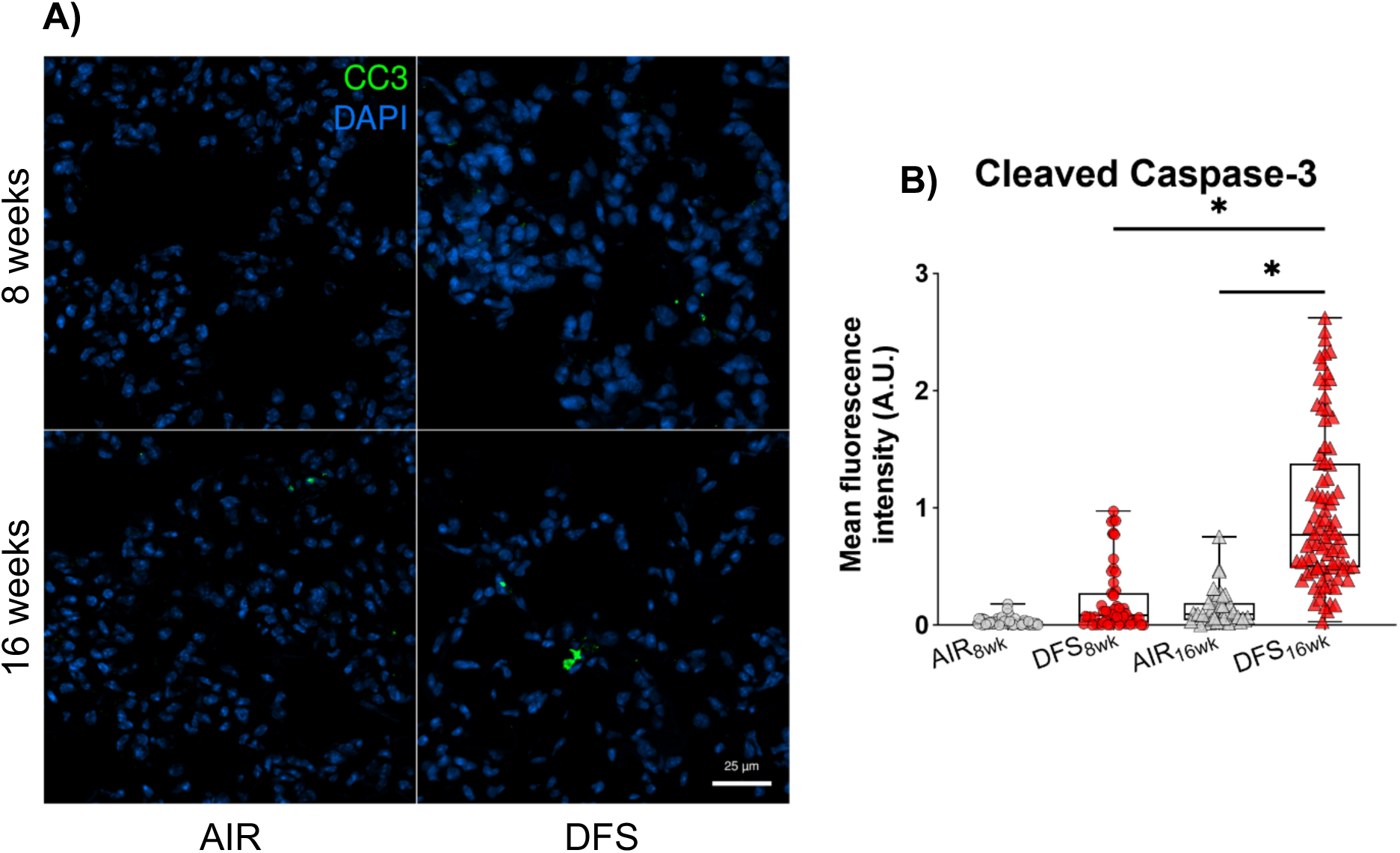
Immunofluorescence analysis of cleaved caspase-3 (CC3) in the lungs’ parenchyma following DFS exposure. A) Representative confocal images of lung sections stained for cleaved caspase-3 (CC3, green) and nuclei (DAPI, blue) from AIR_8wk_, DFS_8wk_, AIR_16wk_, and DFS_16wk_ groups. Late-stage apoptotic cells were identified by distinct nuclear condensation and fragmentation, along with nuclear co-localization of CC-3, which were diffusely present in the DFS_16wk_ group. B) Quantification of the total CC-3 mean immunofluorescence intensity in the alveolar region, presented in arbitrary units (A.U.). A significant increase in CC-3 signal was observed in DFS_16wk_ compared to AIR_16wk_, indicating an apoptotic burden with prolonged smoke exposure. All images were captured at 40x magnification and under the same acquisition parameters. Scale bar = 25 μm. Individual data points represent mean fluorescence intensity per image. Box plots show the median and interquartile range, with whiskers indicating minimum and maximum values (N = 4 mice/group; multiple cross-sections and parenchymal regions per lung). Immunofluorescence data were analyzed using mixed-effects models accounting for within-mouse clustering (ICC), with pairwise comparisons adjusted using FDR; significance was defined as FDR-adjusted p < 0.1 (*).

**Fig. 6.**
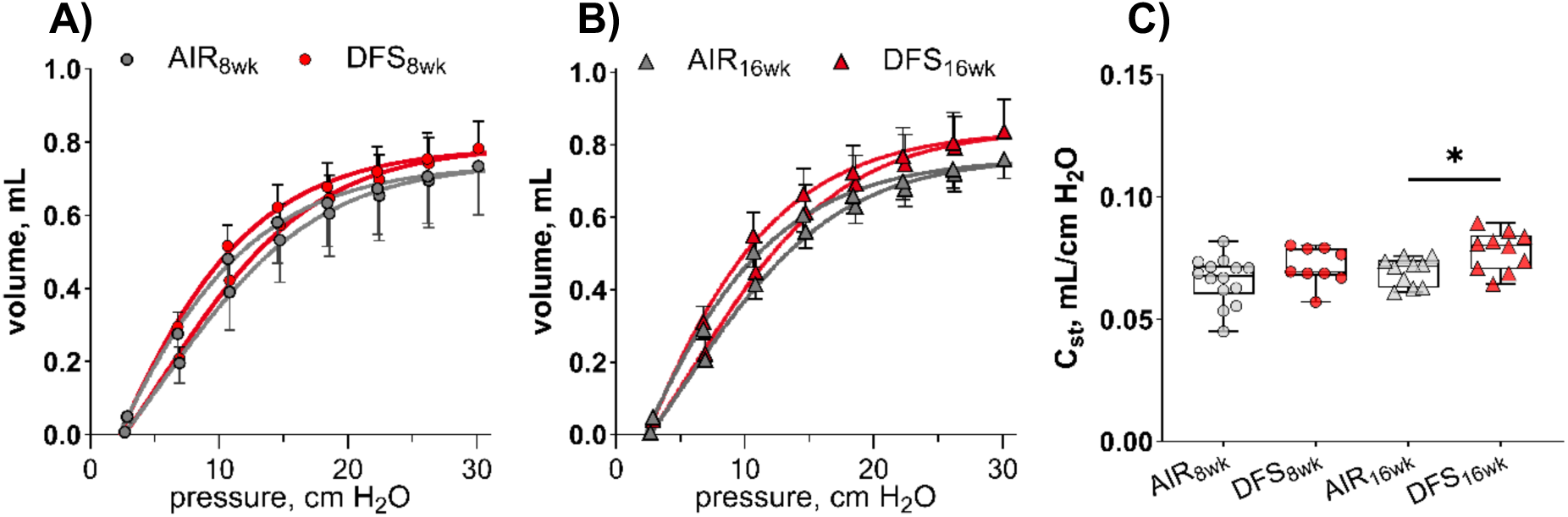
Progressive alteration of pressure-volume (P-V) curves and static compliance following DFS exposure. P-V curves exhibited an upward shift and static compliance increased significantly after 16 weeks of DFS exposure, indicating reduced lung elastic recoil. P-V curves are shown in A) 8 weeks (AIR = 14 compared to DFS= 9 mice), and B) 16 weeks (AIR = 10 compared to DFS = 10 mice). C) Quasi-static compliance (C_st_) was significantly elevated in DFS_16wk_ compared to AIR_16wk_. Group differences were assessed using one-way ANOVA with FDR-adjusted p-values. Statistical significance between experimental groups was defined for FDR-adjusted p < 0.1 (*).

Apoptosis is also implicated in emphysema pathogenesis, even in the absence of inflammation (Kasahara et al. 2000; Aoshiba et al. 2003; Tang et al. 2004; Petrache et al. 2005; Demedts et al. 2006; Fehrenbach 2007), which has been observed in woodsmoke (Ramos et al. 2009) and cigarette smoke (Fehrenbach 2007) exposures. Here, cleaved caspase-3 (CC3, Fig.5) and its upstream marker cleaved caspase-9 (CC-9, Supplementary Material, Fig. S3), canonical markers of apoptosis, showed a progressive increase from 8 to 16 weeks of DFS exposure, mirroring the trend observed in the number of alveolar blunts (Fig. 3G) and static mechanics (Fig. 4). In the absence of cell survival pathways such as NF-κB and EMT, this structural deterioration may enhance anoikis, a form of programmed cell death due to loss of cell-matrix adhesion, thereby contributing to the increased epithelial permeability observed following 16 weeks of exposure (McCawley and Matrisian 2001; Demedts et al. 2006), a process previously described in mouse models of emphysema (Aoshiba et al. 2003; Demedts et al. 2006). Alternatively or concurrently, increased apoptosis may reflect an impaired efferocytosis, potentially resulting from insufficient SP-D mediated stimulation of macrophage clearance or from neutrophil elastase mediated cleavage of macrophage phosphatidylserine receptors (Demedts et al. 2006). This might explain the discordance between SP-D and CC-3 after 8 and 16 weeks of DFS exposure.

An unexpected observation from our study is that early proteolytic response and structural changes preceded a measurable functional decline. It was not until 16 weeks of exposure that we detected measurable changes in static compliance and pressure-volume curves of the lung, corresponding with exacerbated airspace enlargement and alveolar blunting. This structure-function mismatch echoes previous findings in murine woodsmoke exposure (Tesfaigzi et al. 2002), cigarette smoke (Wright and Churg 1990; Guerassimov et al. 2004; Foronjy et al. 2005), and emphysema models induced by elastase (Anciães et al. 2011). In the study by Guerassimov *et al*., Pallid and AKR/J mouse strains, which developed more diffuse emphysema in response to cigarette smoke exposure, exhibited the greatest increases in L_m_ and the most pronounced declines in lung compliance (Guerassimov et al. 2004). In contrast, the C57BL/6 strain showed emphysematous changes primarily localized to the alveolar ducts, with comparatively preserved lung mechanics. Later echoed by Anciães et *al.*, these studies emphasize that a critical threshold of alveolar destruction, reflected by L_m_, must be surpassed before mechanical impairment becomes detectable and sensitive under such respiratory mechanics assessment techniques (Anciães et al. 2011). Robbesom *et al*. contend that the characterization of emphysema should rely primarily on morphological evaluation rather than global lung function measurements, given the consistently weak correlation observed between structure and function (Robbesom et al. 2003). This discrepancy is not unexpected in a disease as spatially heterogeneous as emphysema. Since lung function parameters reflect the integrated performance of the entire organ, they may fail to detect regional destruction captured by histological assessment. Conversely, morphometric analysis remains restricted to the sampled tissue and may not represent the lung’s overall structural integrity. Collectively, this suggests that the distribution and magnitude of alveolar injury, rather than its mere presence, could dictate the onset of functional decline. In scenarios where alveolar destruction remains spatially constrained, global lung recoil may remain functionally intact. By contrast, once airspace enlargement becomes widespread and diffuses, mechanical failure is more likely to manifest and become detectable. This resonates with our structural and functional findings.

Moreover, DFS exposure produced thicker parenchymal septa than AIR controls in both time points, but decreased with DFS exposure. Tissue thickness may counteract functional changes caused by enlarged airspaces. This in turn has likely resulted in only a modest, non-significant increase in C_st_ following 8 weeks of exposure. After 16 weeks of DFS exposure, progressive matrix disruption became evident, as indicated by increased L_m_ (Fig. 3E) and blunting of the alveolar walls (Fig. 3F). Furthermore, the loss of the load-bearing elastin networks, weakened alveolar tethering and interdependence, contribute toward a significant increase in C_st_ and an upward shift of the P-V curves following 16 weeks of DFS exposure.

In conclusion, our murine model recapitulates wildland firefighter occupational exposure scenarios following 7-14 years of service, depending on the length and number of shifts worked. Early changes seen in 8 weeks of DFS exposure could progress to functional decline even after the elastolytic activity wanes. The onset of emphysema-like changes at the longer time point raises concern that occupational exposure may drive respiratory functional decline in this workforce. These findings support the implementation of routine health monitoring of WLFFs and underscore the urgency of implementing preventive protective strategies. The lack of mechanical impairment after 8 weeks of exposure mirrors the asymptomatic period during which firefighters may retain normal lung function, despite accumulating microscopic and molecular damage. Future translational studies should aim to track early-stage biomarkers in firefighters exposed across fire seasons and evaluate their relevance to COPD progression.

## ACKNOLEDGMENTS

We would like to thank Prianka Sarathy at Northeastern University for counting alveolar blunts.

We also thank Tufts Animal Histology Core (AHC) for MOVAT histology services and the Institute for Chemical Imaging of Living Systems (RRID:SCR_022681) at Northeastern University for consultation and imaging support.

## SUPPLEMENTARY MATERIAL

Supplementary material included as submission material.

## Nonstandard Abbreviations and Acronyms

Apoe^-/-^: Apolipoprotein E-deficient (knockout)
CC-3: Cleaved Caspase-3
COPD: Chronic Obstructive Pulmonary Disease
C_st_: Static lung compliance
DFS: Douglas fir smoke
EMT: Epithelial-mesenchymal transition
L_m_: Mean linear intercept
MMP: Matrix metalloproteinase
N-CAD: N-Cadherin
NE: Neutrophil elastase
PM: Particulate matter
P-V: Pressure-volume
SP-D: Surfactant protein-D
WFS: Wildfire smoke
WLFF: Wildland firefighter
ZO-1: Zonula occludens-1

## Methods

### Bronchoalveolar Lavage Fluid (BALF)

Following confirmation of deep anesthesia, lungs were lavaged 3 times (N=5-8 per experimental group) and the recovered BALF was processed using standard protocols (Daubeuf and Frossard 2012). Cytokine concentration was determined in BALF using a multiplex assay (QPlex, Quansys, Biosciences, Logan, UT).

**Fig. S1.**
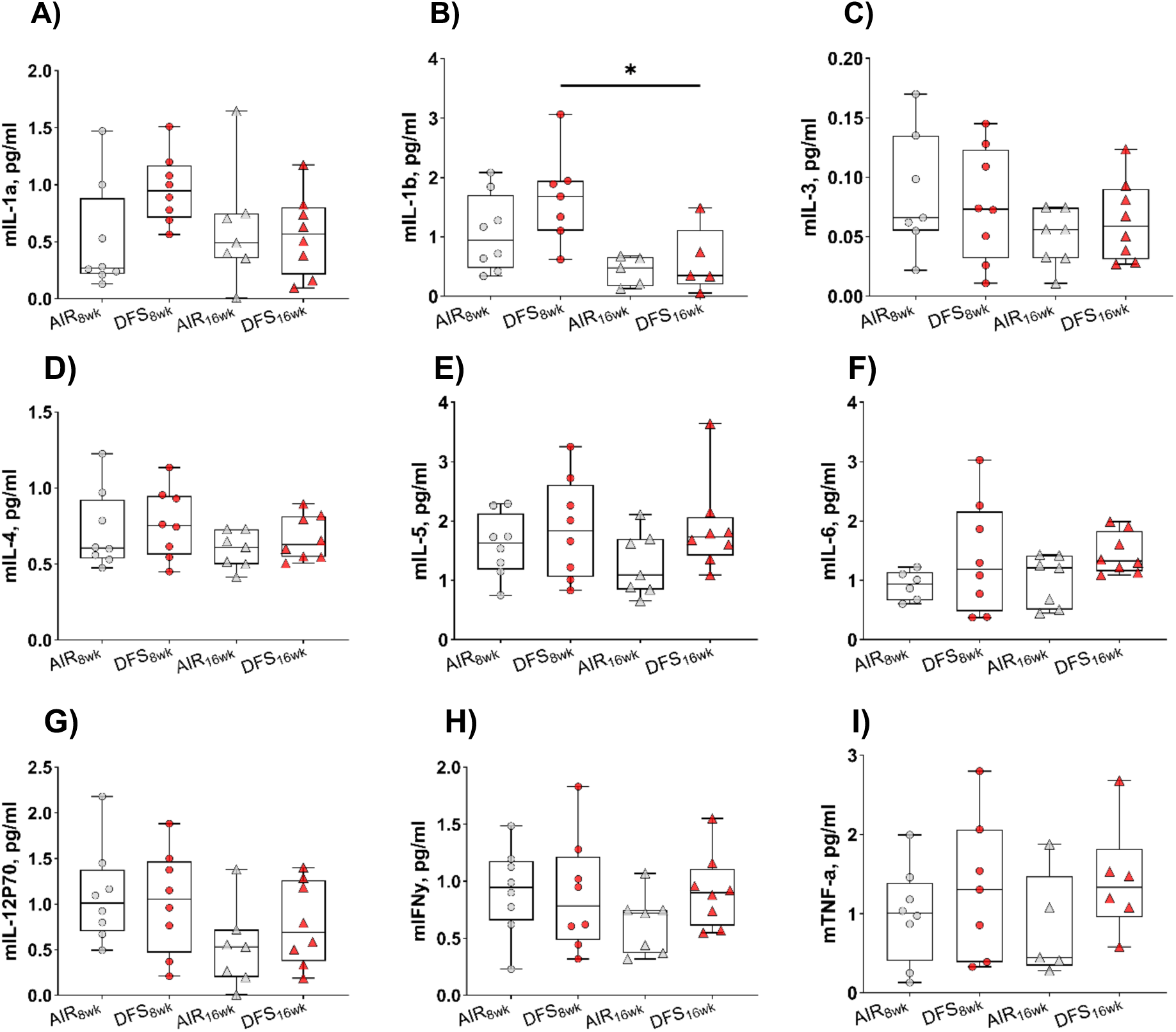
Pro-inflammatory cytokine levels in bronchoalveolar lavage fluid following DFS exposure. Concentrations of pro-inflammatory cytokines in BALF were measured using a multiplex assay from mice exposed to filtered air (AIR) or Douglas fir smoke (DFS) for 8 or 16 weeks. Data are shown for (A) IL-1α, (B) IL-1β, (C) IL-3, (D) IL-4, (E) IL-5, (F) IL-6, (G) IL-12p70, (H) IFNγ, and (I) TNFα. A slight increase in IL-1α (FDR = 0.1969) and IL-1β (FDR = 0.1295) levels was observed in the DFS_8wk_ group compared to AIR_8wk_, but did not reach statistical significance. Across all markers, no significant elevation was detected in DFS-exposed groups relative to their air-exposed counterparts at either time point. Box plots show the median and interquartile range, with whiskers indicating minimum and maximum values, and individual data points representing individual samples. Statistical comparisons were conducted using one-way ANOVA with FDR adjustment. Significance was considered for FDR-adjusted p < 0.1 (*).

**Fig. S2.**
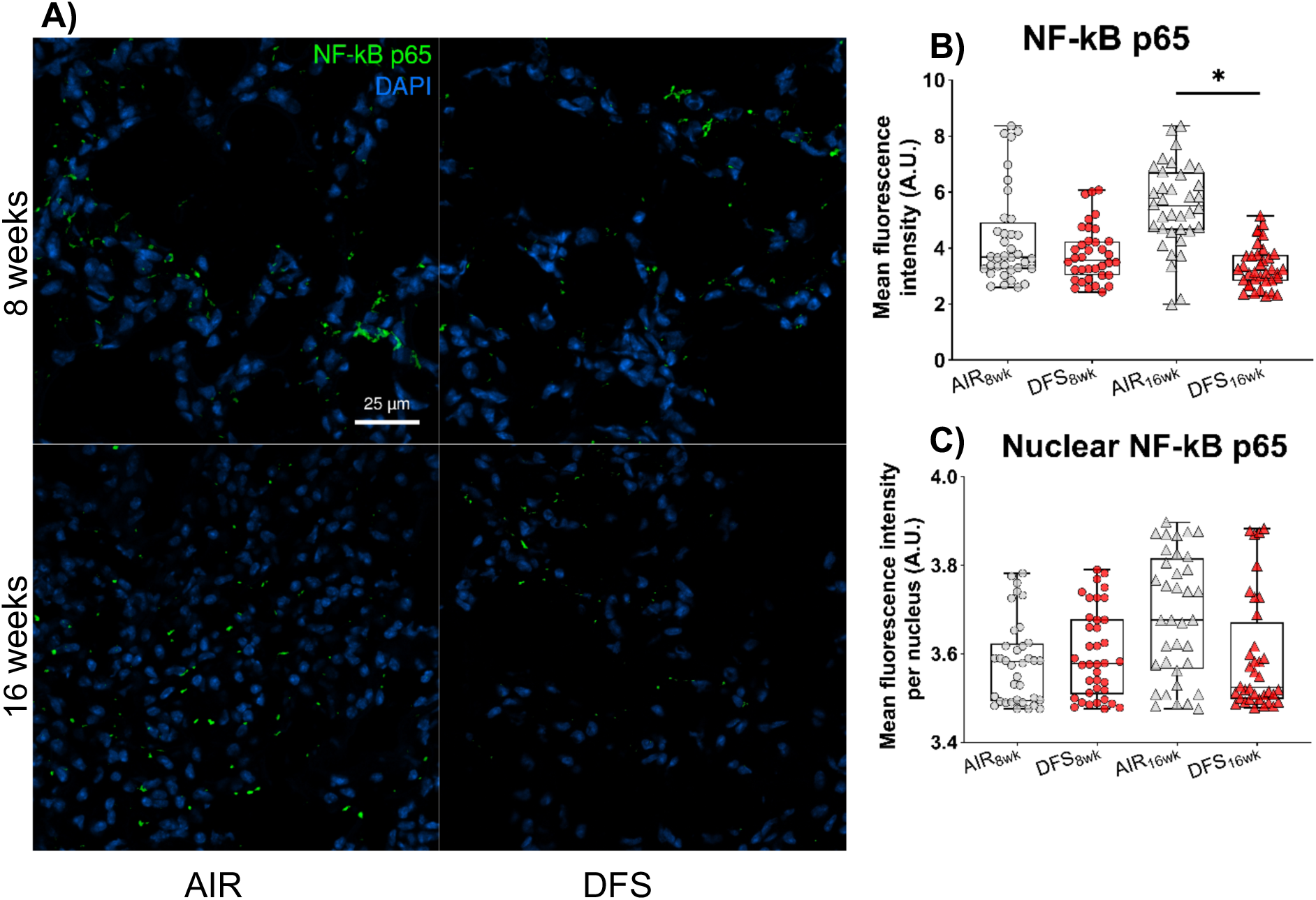
NF-κB p65 immunofluorescence and nuclear translocation in lung parenchyma following DFS exposure. (A) Representative confocal images showing NF-κB p65 (green) and nuclear DAPI staining (blue) in lung sections from AIR_8wk_, DFS_8wk_, AIR_16wk_, and DFS_16wk_ groups. (B) Quantification of total NF-κB p65 fluorescence intensity across treatment groups. Total NF-κB p65 signal was slightly lower in DFS_16wk_ compared to AIR_16wk_ (P = 0.0945). (C) Mean nuclear fluorescence intensity per cell nucleus was used to quantify NF-κB p65 nuclear translocation. No significant changes were observed in any of the groups. At the time points assessed, NF-κB was not an active transcription factor. All images were acquired at 40x magnification and under the same acquisition parameters. Scale bar = 25 μm. Individual data points represent mean fluorescence intensity per surface area. Box plots show the median and interquartile range, with whiskers indicating minimum and maximum values (N = 4 mice/group; multiple cross-sections and parenchymal regions per lung). Statistical comparisons were conducted using one-way ANOVA with FDR adjustment. Significance was considered for FDR-adjusted p < 0.1 (*).

**Fig. S3.**
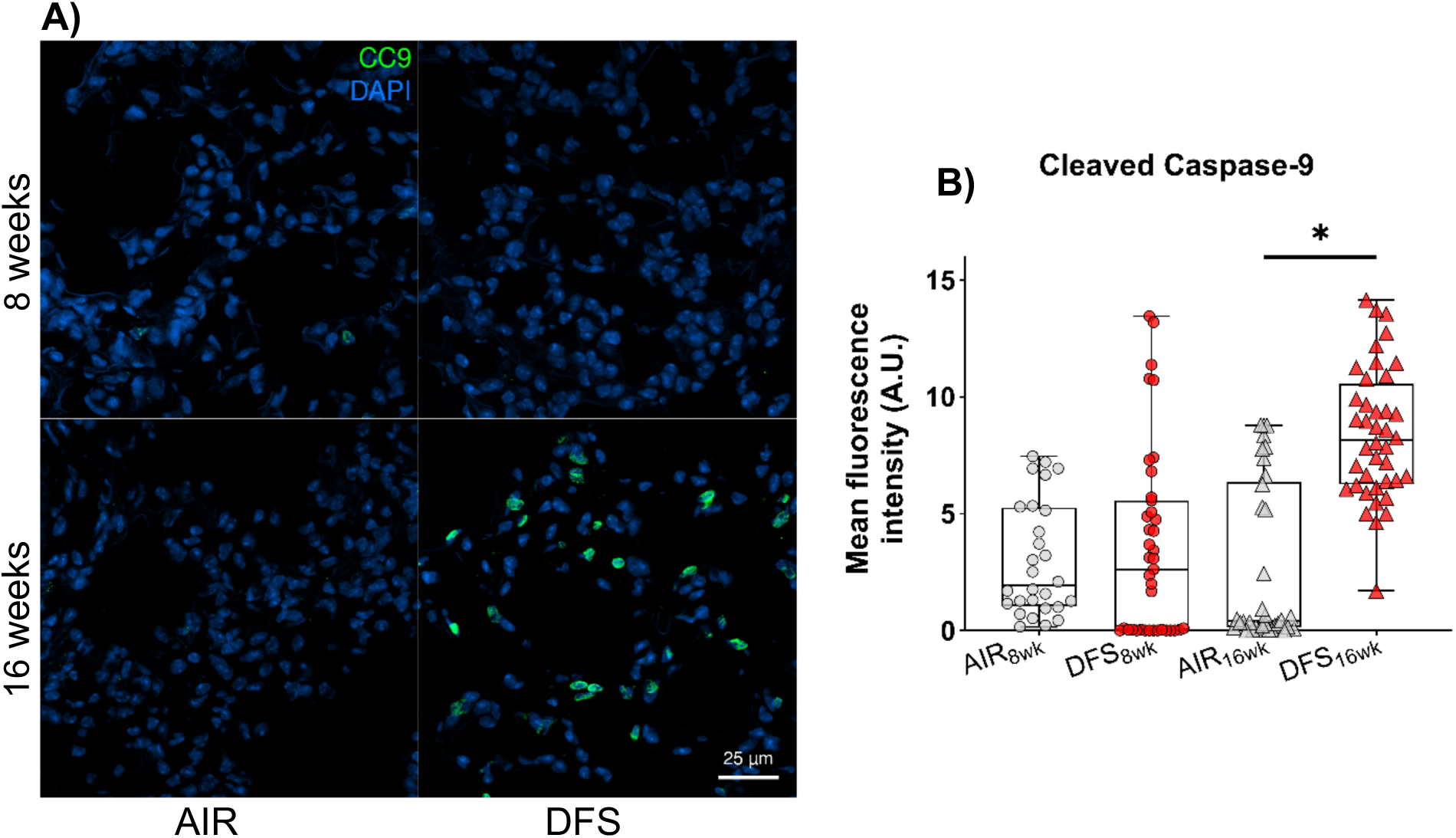
Cleaved caspase-9 immunofluorescence in lung parenchyma following DFS exposure. (A) Representative confocal images showing cleaved caspase-9 (CC-9, green) and nuclear DAPI staining (blue) in lung sections from AIR_8wk_, DFS_8wk_, AIR_16wk_, and DFS_16wk_ groups. Prominent CC-9 staining was observed in DFS_16wk_ lungs (B) Quantification of CC-9 mean fluorescence intensity revealed a significant increase in DFS_16wk_ lungs compared to AIR_16wk_. All images were acquired at 40x magnification and under the same acquisition parameters. Scale bar = 25 μm. Individual data points represent mean fluorescence intensity per image. Box plots show the median and interquartile range, with whiskers indicating minimum and maximum values (N = 3 mice/group; multiple cross-sections and parenchymal regions per lung). Statistical comparisons were conducted using one-way ANOVA with FDR adjustment. Significance was considered for FDR-adjusted p < 0.1 (*).

